# Speech-In-Noise Comprehension is Improved When Viewing a Deep-Neural-Network-Generated Talking Face

**DOI:** 10.1101/2022.07.01.497610

**Authors:** Tong Shan, Chenliang Xu, Zhiyao Duan, Ross K. Maddox

## Abstract

Listening in a noisy environment is challenging, but many previous studies have demonstrated that comprehension of speech can be substantially improved by looking at the talker’s face. We recently developed a deep neural network (DNN) based system that generates movies of a talking face from speech audio and a single face image. In this study, we aimed to quantify the benefits that such a system can bring to speech comprehension, especially in noise. The target speech audio was masked with signal to noise ratios of −9, −6, −3, and 0 dB and was presented to subjects in three audio-visual (AV) stimulus conditions: 1) synthesized AV: audio with the synthesized talking face movie; 2) natural AV: audio with the original movie from the corpus; and 3) audio-only: audio with a static image of the talker. Subjects were asked to type the sentences they heard in each trial and keyword recognition was quantified for each condition. Overall, performance in the synthesized AV condition fell approximately halfway between the other two conditions, showing a marked improvement over the audio-only control but still falling short of the natural AV condition. Every subject showed some benefit from the synthetic AV stimulus. The results of this study support the idea that a DNN-based model that generates a talking face from speech audio can meaningfully enhance comprehension in noisy environments, and has the potential to be used as a “visual hearing aid.”

## Introduction

Speech in quiet environments can usually be easily understood. However, listening effort is increased and comprehension eventually suffers when a speech signal of interest is degraded, either by the presence of additional talkers and other environmental noise or hearing loss, with the combination presenting particular difficulty. Successful listening under such circumstances, often described as the Cocktail Party Problem (Cherry, 1953), involves segregating sound sources and selectively attending the target (McDermott, 2009; Shinn-Cunningham & Best, 2008).

Natural speech, however, is not only an acoustic signal. Listening in noisy situations is aided by a number of factors, principal among them being visual speech cues provided by viewing the talker’s face (Bernstein & Grant, 2009; Erber, 1969; Sumby & Pollack, 1954). However, there are numerous listening situations in which visual speech cues are not available, such as phone calls and podcasts. The COVID-19 pandemic has introduced additional challenges to audio-visual speech perception, with the advent of widespread mask-wearing and video-conferencing in which often one or several participants do not turn on their cameras.

As a result of these challenging listening scenarios, there is interest in how artificial visual cues may aid listening. Recent work has suggested that a temporally coherent non-speech stimulus offers little or no aid in segregating concurrent acoustic stimuli. Strand J et al (2020) used a modulated visual circle presented coherently with speech and found that the circle did not help reducing effort or improving subjects’ speech comprehension. A similar study (Yuan, Lleo, Daniel, White, & Oh, 2021) used a 3-dimentional modulating sphere rather than a 2D circle and found a small improvement which was significant only at a signal-to-noise ratio (SNR) of −1dB. Results have been uninspiring for other modalities as well— Riecke et al (2019) presented subjects with tactile stimuli which were coherent with the speech audio envelope. Although they observed an enhanced cortical tracking of the speech in EEG, there was no improvement in word recognition performance.

In contrast to simple artificial stimuli, the audio-visual benefit of natural speech stimuli is substantial (Bernstein & Grant, 2009; Erber, 1969; Sumby & Pollack, 1954), as well as being robust to asynchrony with the acoustic speech (Grant & Greenberg, 2001) and degradation in spatial frequency (Munhall, Kroos, Jozan, & Vatikiotis-Bateson, 2004).Thus, in listening situations where comprehension would benefit from visual face cues, but none are present, creating a synthetic face that matches the acoustic signal holds promise for assisting listening.

Deep neural networks (DNN) provide an exciting opportunity for synthetically generated visual listening aids but efforts to create such systems predate DNNs’ widespread use. For example, statistical machine learning (ML) methods such as Hidden Markov models (HMM) have been used previously to generate moving mouth animations from either text or speech audio (Al Moubayed, De Smet, & Van Hamme, 2008; Hofer, Yamagishi, & Shimodaira, 2008; Masuko, Kobayashi, Tamura, Masubuchi, & Tokuda, 1998; Schabus, Pucher, & Hofer, 2013; Tamura, Masuko, Kobayashi, & Tokuda, 1998). Other ML methods such as QR factorization (Lucero, Baigorri, & Munhall, 2006) and artificial neural networks (ANN) (Massaro, Beskow, Cohen, Fry, & Rodgriguez, 1999) have also been used for similar purposes. A data-driven talking head product named Baldi (Massaro & Palmer Jr, 1998) used the output of a phoneme recognizer to animate a three-dimensional avatar head. The system was originally designed for language learning and has shown benefit for word recognition for children and multilingual learning (Ouni, Cohen, & Massaro, 2005). The SynFace project (Beskow, Granström, & Spens, 2002; Beskow, Karlsson, Kewley, & Salvi, 2004) was designed as a telephone aid before the advent of smartphones. SynFace used recurrent neural networks (RNN) mixed with HMMs to animate the face, tongue, and teeth of an avatar from acoustical input signal. Improvement was shown in a word-recognition task with SYNFACE added to audio compared with an audio-only condition (Salvi, Beskow, Al Moubayed, & Granström, 2009). The system offered some improvement in listeners with hearing loss, but it was highly variable across listeners.

In recent years, deep learning methods have seen wide adoption in audio-visual synthesis. There have been many studies that use DNN approaches to build audio-to-visual models, aiming to help people with speech intelligence. Some of the studies predicted the canonical face animation by predicting the facial landmarks with long short-term memory (LSTM) network (Eskimez, Maddox, Xu, & Duan, 2018). Chen et al proposed a two-stage system that first converts the speech audio to face landmarks and then maps the landmarks to a reference face image (Chen, Li, Maddox, Duan, & Xu, 2018; Chen, Maddox, Duan, & Xu, 2019). Others focused on generating realistic mouth or facial movement video using convolutional neural networks (CNN) (Jamaludin, Chung, & Zisserman, 2019), LSTM (Pham, Wang, & Pavlovic, 2017; Suwajanakorn, Seitz, & Kemelmacher-Shlizerman, 2017), recurrent adversarial networks (Song, Zhu, Li, Wang, & Qi, 2018), or generative adversarial networks (GAN) (Vougioukas, Petridis, & Pantic, 2019).

Although there are numerous published audio-to-visual DNN models, they generally only assess how realistic the generated face animations are based on quantitative metrics or subjective quality ratings. Little has been done to investigate the comprehension benefit for human listeners especially in difficult listening environments. In one recent exception, Varano et al. (2022) showed a GAN model (Vougioukas, Petridis, & Pantic, 2020) that can improve speech comprehension in noisy environments.

The new DNN model we used in this study (Eskimez, Maddox, Xu, & Duan, 2020) is able to generate realistic talking face videos from pure acoustical signals and a single seed image of a face. The face can be anyone and does not necessarily need be the person who speaks the audio. The model is also designed for real-time processing with very little delay, making it potentially useful for the purpose of communicating over telephone or online meetings with no camera feed. In this study, we investigated if this model specific offers improved speech intelligibility in the presence of background noise. The performance of speech intelligence with three audio-visual (AV) conditions across four different SNRs was compared. We found generally that speech in noise paired with the model-synthesized talking face video showed significantly better speech comprehension than the audio-only control condition, especially in worse SNRs.

## Materials and Methods

In this study participants performed speech-in-noise comprehension tasks in three audio-visual (AV) conditions: 1) Audio-only, 2) Synthesized AV and 3) Natural AV. The target audio was drawn from an open set corpus and was masked with speech from two other talkers at four signal-to-noise ratios (SNRs) in each condition. Participants were asked to replicate the sentence they heard by typing it at a prompt, and the performance was scored based on the percent of keywords correctly reported.

### Subjects

The experiment was conducted under the protocol approved by University of Rochester Research Subjects Review Board. Every subject gave written informed consent before the experiment. Subjects were paid for the time in the lab.

Ten subjects (four male and six female) aged between 19 to 40 (mean 27.5) were included in this study. Subjects underwent an audiometric screen and were confirmed to have audiometric thresholds of 20 dB HL or better at octave frequencies from 500 Hz to 8000 Hz before the experiment. All subjects reported normal or corrected-to-normal vision and English as their primary language. No subjects were excluded.

### Stimulus Materials

The end-to-end DNN-based model developed by Eskimez et al. (2020) was used to generate the synthesized face video. The model takes one audio file and a single image of face as the input. The output video is a cropped talking face with 25 frames / s and 128 by 128 pixels that includes only the facial area of the talker (as shown in **Figure 1**). Please refer to the Eskimez et al (2020) article for detailed description about the model. The model depends only on 136 ms of audio from the “future,” and thus can be run in near real-time with a 136 ms lag. It can be modified to have a shorter lag at the cost of some quality.

**Figure 1.**
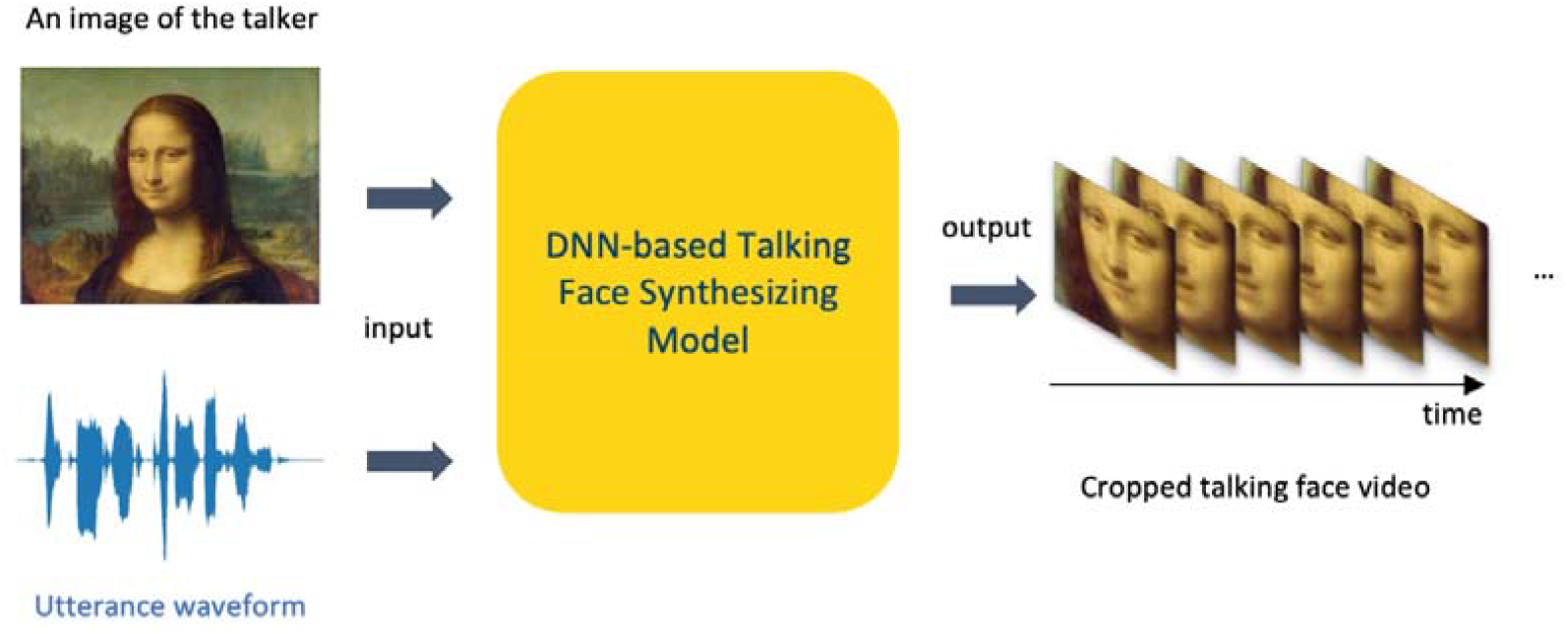
DNN-based model that generates synthesized talking face movie from a pure audio and a single image of face.

The stimuli in this study were selected from the Speech Test Video Corpus (STeVi) (Sensimetrics, 2014), which contains audio and videos of four native English narrators (two male and two female) speaking sentences. Each utterance includes three to five keywords. For example, one utterance is “his PLANS MEANT TAKING a BIG RISK”, where the uppercase words are predetermined to be the keywords.

To study the synthetic video’s effect on speech comprehension in noise, we mixed the target audio from the STeVi corpus with two streams of audio maskers with SNRs of −9, −6, −3, and 0 dB. The maskers were selected from two audiobooks, *The Alchemyst* read by a male narrator (Scott, 2007) and *A Wrinkle in Time* read by a female narrator (L’Engle, 2012). The audiobooks were processed by automatically cutting out the silence period that was longer than 0.5 s using tools developed by our lab previously (Maddox & Lee, 2018; Polonenko & Maddox, 2021). The SNR refers to the ratio of the intensity of the target audio to the intensity of the combined maskers. Each masker was presented at 60 dB SPL, meaning the level of the summed maskers was 63 dB on every trial. The level of the target ranged from 51 to 63 dB SPL, corresponding to SNRs from −9 to 0 dB.

### AV Conditions

The three AV conditions and the four SNRs of noise conditions were randomly paired and presented to the subjects (see **Figure 2**). The three AV conditions were determined entirely by the visual stimulus—the only acoustic manipulation in the experiment was SNR.

**Figure 2.**
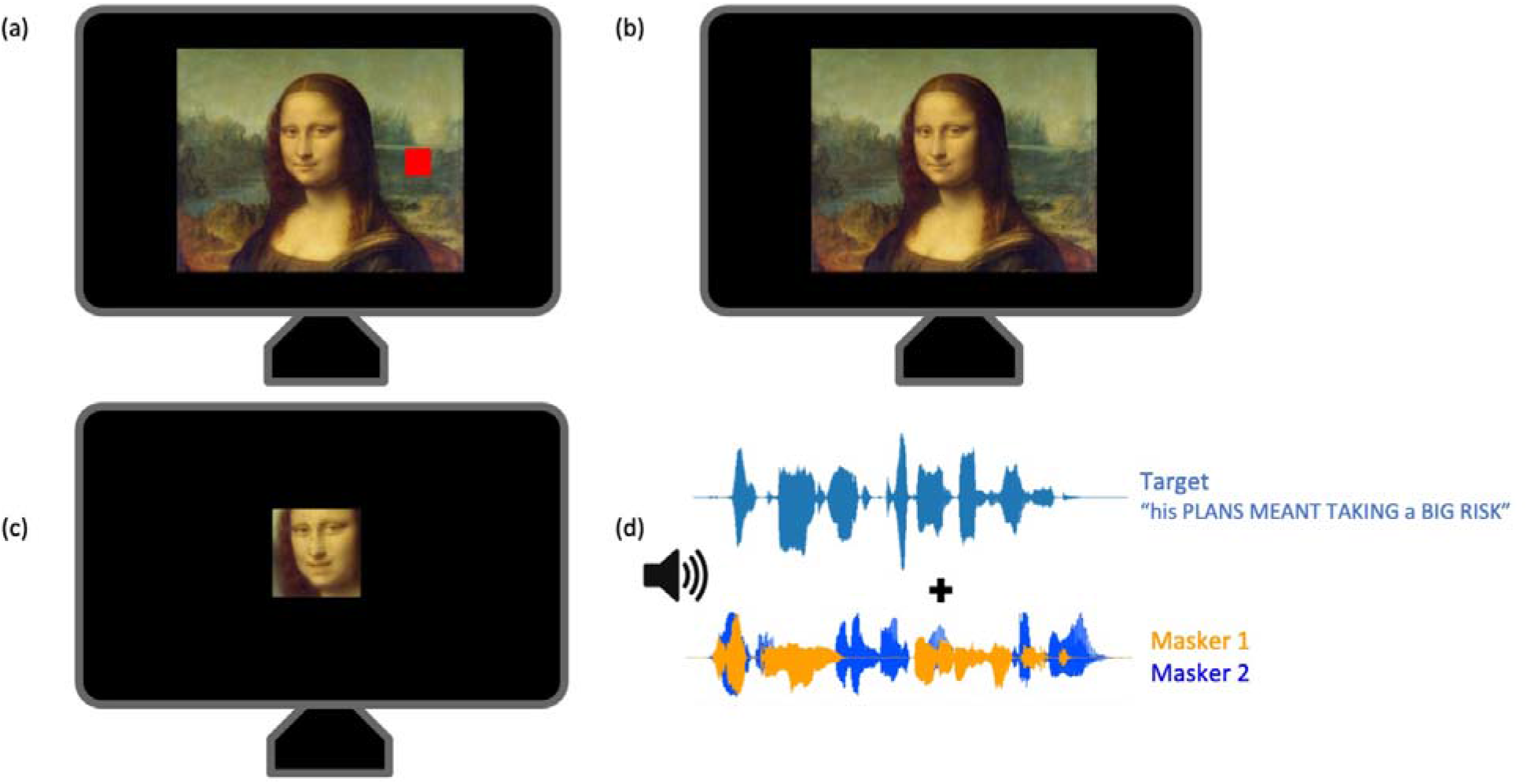
Visual and audio stimuli presented to subjects. (a) Audio-only condition: A static image of the narrator was shown to indicate the narrator’s identity and a small red square on the right was also shown to inform the audio-only condition without any moving video followed. (b) Natural AV condition: The original video from the corpus recording was presented. (c) Synthesized AV condition: The talking face video synthesized using the target speech was presented. (d) Audio signal: target sentences (upper) and two streams of maskers (lower).

1. Audio-only condition: The audio from the corpus that mixed with maskers were presented. The visual stimulus was a static image of the narrator to indicate the narrator’s identity. A small red square on the right was also shown to inform the audio-only condition without any moving video followed (**Figure 2a**). The image was a screenshot taken from the original video of the corpus.
2. Natural AV: The original video from the corpus recording was presented in the center of the screen with 29.97 frames / s, its native framerate (**Figure 2b**). The target audio was also recorded within the video.
3. Synthesized AV: The talking face video synthesized using the target speech was presented. An image of the actual talker was used as the seed, so that the talker identity matched in all trials. Note that none of the audio or video used in this experiment was present in the model’s training set (Eskimez et al., 2020). In this condition, the videos were played at 25 frames / s which is the default output of the model (**Figure 2c**).

The model synthesizes only the narrator’s face. Therefore, to match the size of faces in all conditions, we increased the synthesized video (whose native resolution is only 128 by 128 pixels). We also reduced the size of original video from the corpus respectively, so that the matched synthesized face would not look like pixelated. The angle of the faces subtended is around 6.2° from subjects’ eyes.

Each trial began with a 2 second pause on the first frame of the video to help the subject figure out which AV condition and which voice to listen to for that trial.

### Procedure

Subjects were seated in a sound-isolating booth in front of a 24-inch BenQ monitor with a refresh rate of 144 Hz and viewing distance of around 60 cm. The audio stimuli were played through Etymotic Research ER-2 insert earphones via and RME Babyface Pro sound card. A standard keyboard was given to the subjects to type their response in the experiment.

An acclimation session was first presented for the subjects to get familiar with the voice and match the voices and the faces of the four narrators from the corpus. Three nonsense utterances were played that said by each narrator. Fake names were given to each narrator to facilitate remembering them (“Anna”, “Bella”, “Charles” and “David”). This step was necessary so that subjects could distinguish the target voice from the masker talkers in low SNR conditions.

A 12-trial training session next allowed subjects to become familiar with each AV condition and understand how to respond correctly. Subjects were asked to type in the sentence they heard word by word after each trial. For each of the three AV conditions, there were two trials of clean target audio (without any background noise) and then two trials of audio in noise with the SNRs of 0 and −3 dB. The accuracy for the training session was estimated automatically after each trial. The responses in each trial were considered as correct if the sequence of letters of the entered keywords were no less than 80% correct, and the correctness was presented to the subjects immediately after they submitted their responses (N.B.: automatic evaluation was used only for training; manual evaluation is described in the next section). This ratio was calculated by the SequenceMatcher class of the difflib python library, as in Fiscella, Cappelloni, and Maddox (2022). To pass the training session, subjects needed to report all keywords correctly for the clean trials, and at least one keyword for the 0 dB and −3 dB noise conditions. A second chance to pass the training would be given if subjects could not pass on their first try.

Subjects could proceed to the actual experiment after passing the training session. The 12 stimulus conditions (three AV conditions paired with 4 SNRs) were presented in a completely random order. Each trial presented a unique target sentence. There were 15 trials within each combination of AV and SNR condition. A total of 180 trials were presented to the subjects. Subjects were given breaks with a minimum duration of 30 seconds at 25, 50, and 75 percent completion of the experiment. Responses were recorded and then exported to an excel file for later evaluation.

### Data Analysis

Subjects’ performance was scored manually by counting the correct keywords based on Smayda et al.’s (2016) criteria and previously used by our lab (Fiscella et al., 2022). Responses were considered correct if the typos did not change the meaning of the word or if the words were homophones. Note that this manual scoring was performed offline, and the automated scoring described previously was only used in the training session so feedback could be immediate. The percent correct scores from each of the 12 conditions were calculated for each subject and were averaged over subjects.

A generalized linear mixed-effects model was constructed using *glmer* function from the *lme4* (Bates, Mächler, Bolker, & Walker, 2014) package in R. The model was fitted with a random effect on the subjects. The fixed effects included SNR as a continuous variable, the AV condition as a categorical variable, and the interaction of these two variables (see the formula below). The correctness of each word was treated as a binomial (logit) outcome.

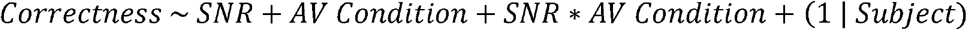

## Results

The generalized linear regression model showed that in the range of SNRs tested, the performance was correlated to the SNR (p < 0.001), with significantly different intercepts for each AV condition. Among the three AV conditions, the order of the performance is Natural AV > Synthesized AV > Audio-only (p<0.001). These results also concluded that as the SNR increases, the performance increases in each AV condition, as expected. There was also an interaction of AV condition and SNR, which reflects the fact that the slope for each AV condition over SNR also differed from each other, with the Audio-only condition showing the largest changes over SNR and Natural AV the smallest (**Table 1**).

**Table 1.** Generalized linear mixed regression model summary. Each row represents the coefficient term with their log odds estimate, the standard error, and the p-values. *p<0.05, **p<0.01, ***p<0.001.

| Variables | Coefficient estimate | Std. Error | p-value |
| --- | --- | --- | --- |
| **Intercept (Audio-only)** | 0.440 | 0.833 | 0.59778 |
| **Synthesized AV** | 0.520 | 0.128 | **$5.17 \times 10^{-5***}$** |
| **Natural AV** | 1.011 | 0.139 | **$3.55 \times 10^{-13***}$** |
| **SNR (Audio-only)** | 0.329 | 0.017 | **$< 2 \times 10^{-16***}$** |
| **Synthesized AV \* SNR** | -0.083 | 0.023 | **$0.00027***$** |
| **Natural AV \* SNR** | -0.136 | 0.024 | **$1.25 \times 10^{-8***}$** |

The average keyword recognition performance in each AV condition and SNR is shown in **Figure 3**. We performed a post hoc pairwise t-test to compare the performance of each AV condition within each SNR condition. The performance of Synthesized AV outperformed Audio-only condition significantly, and closed half of the gap between Natural AV and Audio-only conditions at low SNRs (−9, −6, and −3 dB). Improvements of 20.2%, 21.9%, and 16.3% for Synthesized AV versus the Audio-only condition at −9, −6, and −3 dB of SNR were found, respectively (p<0.05, Holm-Bonferroni corrected). The Natural AV outperformed the Audio-only condition significantly at all of the SNRs, as expected. There was not a significant difference between Synthesized AV and Natural AV at the higher SNRs of −3 and 0 dB.

**Figure 3.**
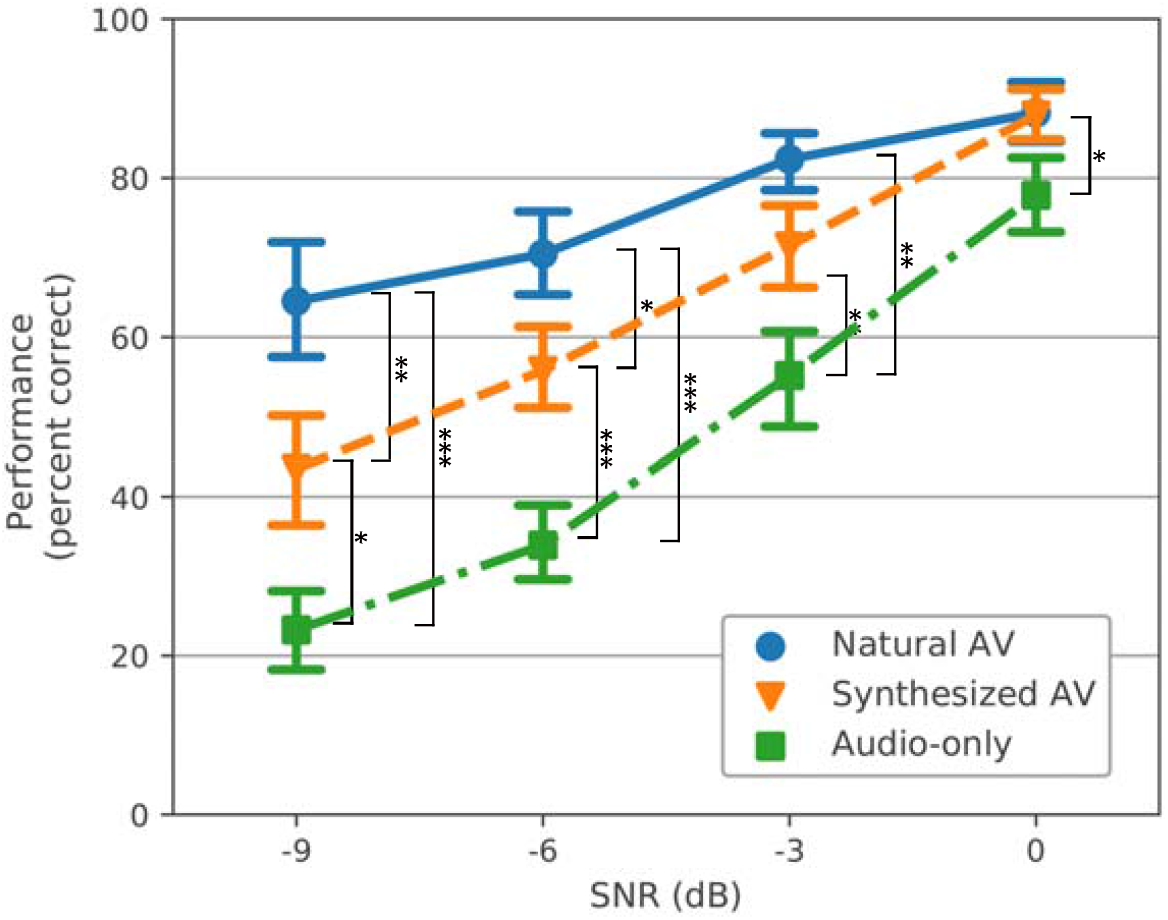
**The grand mean performance (percent correct) across subjects of each AV condition (colored lines) and SNR (bars represent** ±**1 SEM). Brackets indicate significant differences between AV conditions at each SNR (*p<0.05, **p<0.01, ***p<0.001). The bracket at 0 dB indicates a significant difference only between Natural AV and Audio-only.**

Plotting individual subjects’ performance reveals that they all received some benefit from the Synthesized AV condition compared to Audio-only, and some subjects even showed higher accuracies in the Synthesized AV condition than the Natural AV at high SNRs (**Figure 4**). The distribution of the performance differences between Synthesized AV and Audio-only from all SNR conditions across all subjects are shown in **Figure 5**. The kernel density estimate (KDE) curve is highest around a 20 percent improvement, and the large majority of points are positive (i.e., to the right of the line at 0 percent).

**Figure 4.**
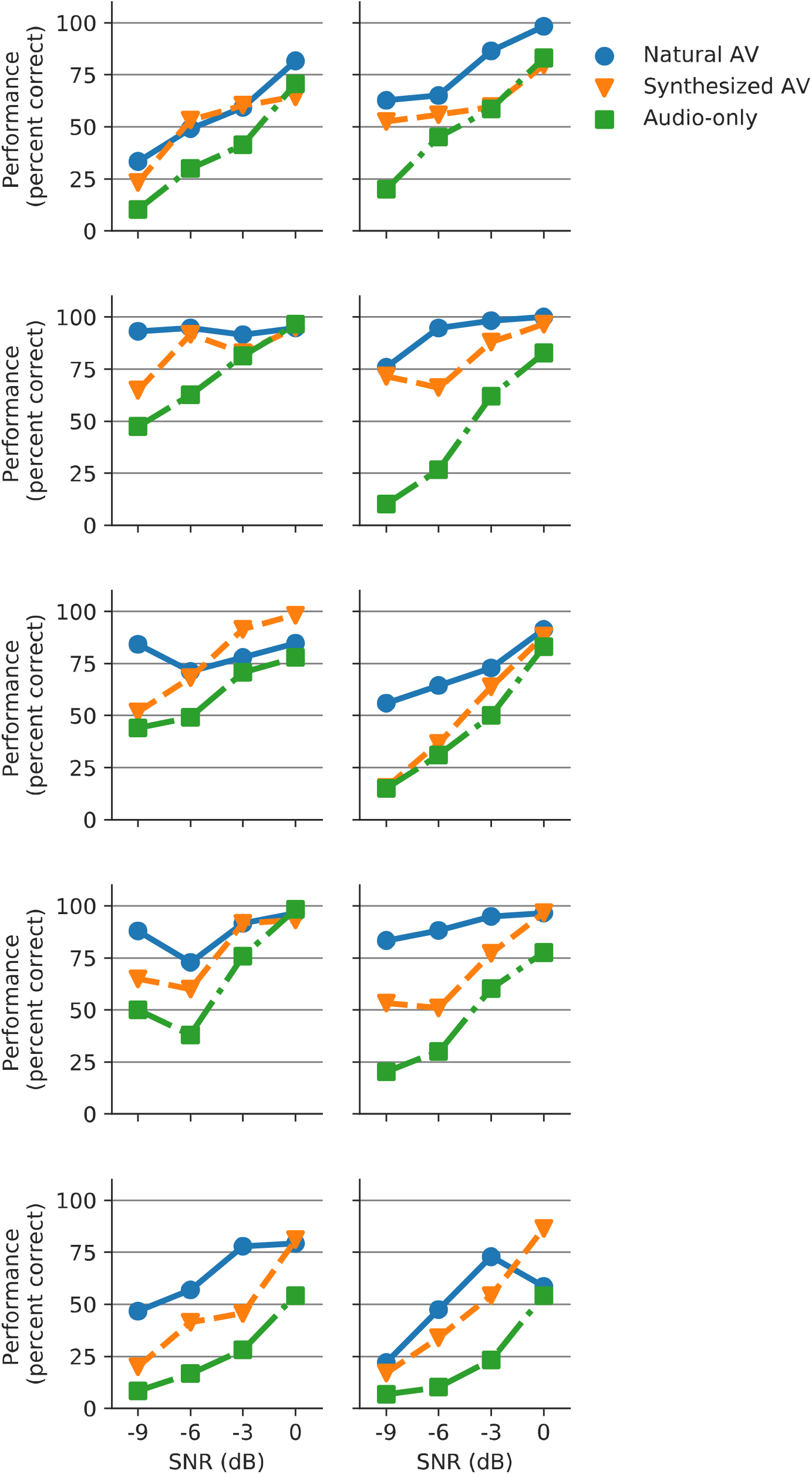
Individual performances (percent correct) from all of the ten subjects.

**Figure 5.**
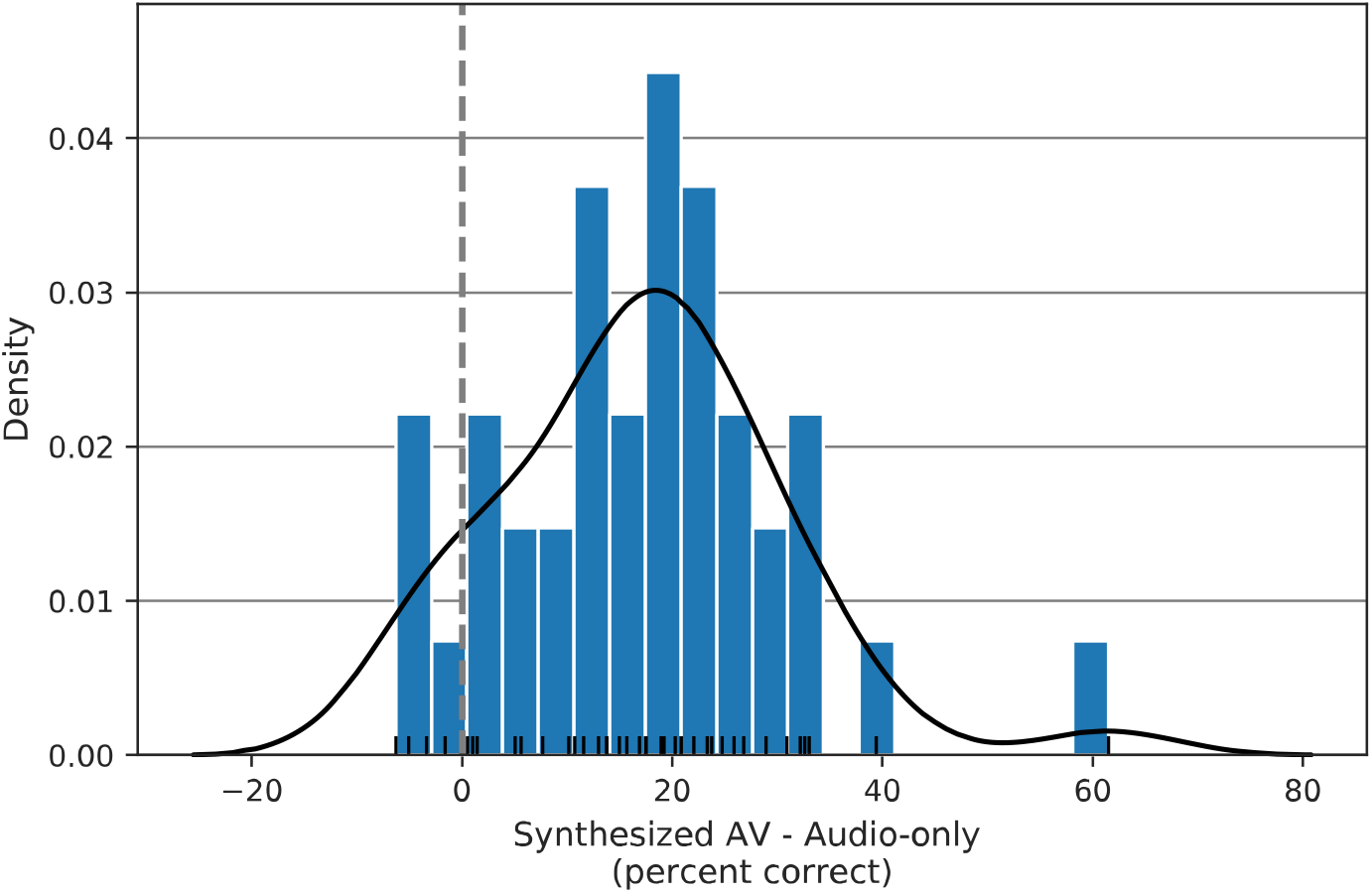
Distribution of the benefit from Synthesized AV. The blue bars are the histogram of the performance (percent correct) difference between the synthesized AV and audio-only condition, taken across all subjects for all SNRs (10 subject * 4 SNR = 40 points). The black curve is the kernel density estimate (KDE) and the short black straight lines indicate individual values. All values to the right of 0 indicate a performance benefit of the synthetic face over the Audio-only control.

## Discussion

Listeners’ performance on the experimental task showed that the synthesized face significantly improves speech comprehension compared to when only acoustical signals are present. The benefit was greatest when listening in low SNRs, with performance in the synthetic face condition falling approximately halfway between the audio-only and natural AV conditions. In the highest SNR (0 dB), performance with the synthesized face was comparable with that of the natural AV condition. Individual results also showed that every subject received some benefit from the model-synthesized face compared to audio-only. The fitted generalized linear model confirmed a significant relationship between the subject performance and the stimulus AV and SNR conditions as well as their interaction.

This study demonstrates that in the absence of a natural visual stimulus, speech comprehension can be enhanced by a synthesized, realistic talking face that is generated purely from the acoustical signal using a DNN-based model. This result is consistent with the recently submitted preprint by Varano et al (2022), where they showed that a GAN model improves speech comprehension in noise both for human and an AI speech recognition system, although they also found the natural face performed better than the model-generated face. That study tested speech comprehension in only one SNR (−8.82 dB) with speech-weighted noise as the background noise. Here, we go beyond that result by testing comprehension in different SNR conditions and using real speech as the background noise.

The synthetic faces in both this study and in Varano et al. (2022) provided a benefit approximately half as large as the natural face did. We believe these benefits stem from realistic faces and especially accurate mouth and articulator shapes. Previous studies using basic visual stimuli such as circles or spheres that dilate coherently with the audio provide no or very small improvements (Strand et al., 2020; Yuan et al., 2021; Yuan, Wayland, & Oh, 2020). In a pilot we ran preceding this study, we used a cartoonish face whose mouth moved coherently but showed little differentiation in its articulation of different sounds. That system also provided no improvement over the audio-only condition. Therefore, we believe that the difference between the improvement offered by the natural and synthetic faces is likely due to small imperfections and inaccuracies yet to be corrected. These imperfections may mean that the synthetic face is 1) not providing some information that a natural face would, 2) providing incorrect information which leads to errors, or 3) interrupting the binding between the audio and visual signals, meaning that some visual information is present but not integrated (or some combination of those factors). New systems that emphasize accuracy of articulation in addition to realism of the face will correct these errors, providing further benefit.

Since the DNN model we employed was designed to run in real time, it has the potential to be considered a “visual hearing aid”. Older systems such as SynFace (Salvi et al., 2009) and Baldi (Massaro & Palmer Jr, 1998) used a phonetic recognizer to first identify a phoneme and then project its articulation to the motion of the 3D head. Different from those systems, our end-to-end model did not include any audio pre- or post-processing or make use of metadata like word- or phoneme-level transcripts during training. It thus was language-agnostic and provides good synchrony. Our model also generates images realistic enough to avoid the “uncanny valley” due to being trained with a GAN approach. The model was also trained with speech in several different noise conditions, meaning the model retains some ability to generate talking faces even if the speech audio is noisy. With these advantages, the developed DNN model may be useful for listening to speech in a wide variety of listening conditions when a talker’s face is not available.

Despite the promising results, some limitations should be noted. First, we recruited only English speakers and presented them with utterances only in English. Although the DNN model was trained on English data, it has the ability to generate talking faces in other languages too. Future work should analyze how well this model works with other languages. Second, in addition to the moving lips and faces that provide the visual speech cues, non-verbal gestures and other spontaneous movements (blinking, for example) also have an impact on normal communication (Cassell, McNeill, & McCullough, 1999; Dargue, Phillips, & Sweller, 2021; Goldin-Meadow, 1999). A model that takes the head movement into account has been developed (Chen et al., 2020) that allows a potential future investigation on what impact of such model can bring in speech comprehension. Emotional expression of the talking face is another influential factor in speech. Changing the emotional expression of the face can lead to changes of the meaning of that utterance (Alpert, Kurtzberg, & Friedhoff, 1963). Another recent DNN-based model is capable of generating talking faces with a categorical emotion input that can be the same as or different from the emotion conveyed in the speech audio (Eskimez, Zhang, & Duan, 2021). It will be interesting to see if such a model can improve speech comprehension more than only neutral faces. Third, we tested the system only in people with typical hearing thresholds. The benefit for people with hearing loss is not known but is of great importance. Successful deployment of a system such as this one will depend on working closely with potential users. With DNN techniques being rapidly developed, face-synthesis models like ours can be (and will need to be) improved, with the eventual goal of providing a benefit comparable to natural faces.

## Acknowledgements

This work was presented at the Association for Research in Otolaryngology Mid Winder Meeting (ARO MWM) 2021 (Shan & Maddox, 2021). Research reported in this publication was supported by the National Institute for Deafness and Other Communication Disorders awarded to RKM (R00DC014288) and Augmented / Virtual Reality pilot grant awarded by the University of Rochester to RKM, CX, and ZD.

## Notes

### Competing Interest Statement

The authors have declared no competing interest.

## Reference

Al Moubayed, S., De Smet, M., & Van Hamme, H. (2008). Lip Synchronization: from Phone Lattice to PCA Eigen-projections using Neural Networks. In Proc. Interspeech 2008 (pp. 2016–2019): ISCA-INST SPEECH COMMUNICATION ASSOC.

Alpert, M., Kurtzberg, R. L., & Friedhoff, A. J. (1963). Transient voice changes associated with emotional stimuli. Archives of General Psychiatry, 8(4), 362–365. doi:https://doi.org/10.1001/archpsyc.1963.01720100052006

Bates, D., Mächler, M., Bolker, B., & Walker, S. (2014). Fitting linear mixed-effects models using lme4. arXiv preprint 1406.5823. doi:https://doi.org/10.48550/arXiv.1406.5823

Bernstein, J. G., & Grant, K. W. (2009). Auditory and auditory-visual intelligibility of speech in fluctuating maskers for normal-hearing and hearing-impaired listeners. The Journal of the Acoustical Society of America, 125(5), 3358–3372. doi:https://doi.org/10.1121/1.3110132

Beskow, J., Granström, B., & Spens, K.-E. (2002). Articulation strength-Readability experiments with a synthetic talking face. In Proceedings of Fonetik 2002 (pp. 97–100): Citeseer.

Beskow, J., Karlsson, I., Kewley, J., & Salvi, G. (2004). Synface–a talking head telephone for the hearing-impaired. In Computers Helping People with Special Needs (pp. 1178–1185): Springer Berlin Heidelberg.

Cassell, J., McNeill, D., & McCullough, K.-E. (1999). Speech-gesture mismatches: Evidence for one underlying representation of linguistic and nonlinguistic information. Pragmatics & cognition, 7(1), 1–34. doi:https://doi.org/10.1075/pc.7.1.03cas

Chen, L., Cui, G., Liu, C., Li, Z., Kou, Z., Xu, Y., & Xu, C. (2020). Talking-Head Generation with Rhythmic Head Motion. In (pp. 35–51). Cham: Springer International Publishing.

Chen, L., Li, Z., Maddox, R. K., Duan, Z., & Xu, C. (2018). Lip movements generation at a glance. In Proceedings of the European Conference on Computer Vision (ECCV) (pp. 520–535).

Chen, L., Maddox, R. K., Duan, Z., & Xu, C. (2019). Hierarchical cross-modal talking face generation with dynamic pixel-wise loss. In Proceedings of the IEEE/CVF Conference on Computer Vision and Pattern Recognition (pp. 7832–7841).

Cherry, E. C. (1953). Some experiments on the recognition of speech, with one and with two ears. The Journal of the acoustical society of America, 25(5), 975–979. doi:https://doi.org/10.1121/1.1907229

Dargue, N., Phillips, M., & Sweller, N. (2021). Filling in the gaps: observing gestures conveying additional information can compensate for missing verbal content. Instructional Science, 49(5), 637–659. doi:https://doi.org/10.1007/s11251-021-09549-2

Erber, N. P. (1969). Interaction of audition and vision in the recognition of oral speech stimuli. Journal of speech and hearing research, 12(2), 423–425. doi:https://doi.org/10.1044/jshr.1202.423

Eskimez, S. E., Maddox, R. K., Xu, C., & Duan, Z. (2018). Generating talking face landmarks from speech. In International Conference on Latent Variable Analysis and Signal Separation (pp. 372–381): Springer.

Eskimez, S. E., Maddox, R. K., Xu, C., & Duan, Z. (2020). End-To-End Generation of Talking Faces from Noisy Speech. In ICASSP 2020-2020 IEEE International Conference on Acoustics, Speech and Signal Processing (ICASSP) (pp. 1948–1952): IEEE.

Eskimez, S. E., Zhang, Y., & Duan, Z. (2021). Speech driven talking face generation from a single image and an emotion condition. IEEE Transactions on Multimedia. doi:https://doi.org/10.1109/TMM.2021.3099900

Fiscella, S., Cappelloni, M. S., & Maddox, R. K. (2022). Independent mechanisms of temporal and linguistic cue correspondence benefiting audiovisual speech processing. Attention, Perception, & Psychophysics, 1–11. doi:https://doi.org/10.3758/s13414-022-02440-3

Goldin-Meadow, S. (1999). The role of gesture in communication and thinking. Trends in cognitive sciences, 3(11), 419–429. doi:https://doi.org/10.1016/S1364-6613(99)01397-2

Grant, K. W., & Greenberg, S. (2001). Speech intelligibility derived from asynchronous processing of auditory-visual information. In AVSP 2001-International Conference on Auditory-Visual Speech Processing.

Hofer, G., Yamagishi, J., & Shimodaira, H. (2008). Speech-driven lip motion generation with a trajectory HMM. In in Proc. Interspeech 2008 (pp. 2314–2317).

Jamaludin, A., Chung, J. S., & Zisserman, A. (2019). You said that?: Synthesising talking faces from audio. International Journal of Computer Vision, 127(11), 1767–1779. doi:https://doi.org/10.1007/s11263-019-01150-y

L’Engle, M. (2012). A Wrinkle in Time. New York: Listening Library.

Lucero, J. C., Baigorri, A. R., & Munhall, K. G. (2006). Data-driven facial animation of speech using a QR factorization algorithm. In Proc. 7th Int. Sem. Speech Prod (pp. 135–142).

Maddox, R. K., & Lee, A. K. (2018). Auditory brainstem responses to continuous natural speech in human listeners. Eneuro, 5(1). doi:https://doi.org/10.1523/ENEURO.0441-17.2018

Massaro, D. W., Beskow, J., Cohen, M. M., Fry, C. L., & Rodgriguez, T. (1999). Picture my voice: Audio to visual speech synthesis using artificial neural networks. AVSP’99-International Conference on Auditory-Visual Speech Processing.

Massaro, D. W., & Palmer Jr, S. E. (1998). Perceiving talking faces: From speech perception to a behavioral principle: Mit Press.

Masuko, T., Kobayashi, T., Tamura, M., Masubuchi, J., & Tokuda, K. (1998). Text-to-visual speech synthesis based on parameter generation from HMM. In Proceedings of the 1998 IEEE International Conference on Acoustics, Speech and Signal Processing, ICASSP’98 (Cat. No. 98CH36181) (Vol. 6, pp. 3745–3748): IEEE.

McDermott, J. H. (2009). The cocktail party problem. Current Biology, 19(22), R1024–R1027. doi:https://doi.org/10.1016/j.cub.2009.09.005

Munhall, K. G., Kroos, C., Jozan, G., & Vatikiotis-Bateson, E. (2004). Spatial frequency requirements for audiovisual speech perception. Perception & Psychophysics, 66(4), 574–583. doi:https://doi.org/10.3758/BF03194902

Ouni, S., Cohen, M. M., & Massaro, D. W. (2005). Training Baldi to be multilingual: A case study for an Arabic Badr. Speech Communication, 45(2), 115–137. doi:https://doi.org/10.1016/j.specom.2004.11.008

Pham, H. X., Wang, Y., & Pavlovic, V. (2017). End-to-end learning for 3d facial animation from raw waveforms of speech. arXiv preprint 1710.00920. doi:https://doi.org/10.48550/arXiv.1710.00920

Polonenko, M. J., & Maddox, R. K. (2021). Exposing distinct subcortical components of the auditory brainstem response evoked by continuous naturalistic speech. Elife, 10, e62329. doi:https://doi.org/10.7554/eLife.62329

Riecke, L., Snipes, S., van Bree, S., Kaas, A., & Hausfeld, L. (2019). Audio-tactile enhancement of cortical speech-envelope tracking. NeuroImage, 202, 116134. doi:https://doi.org/10.1016/j.neuroimage.2019.116134

Salvi, G., Beskow, J., Al Moubayed, S., & Granström, B. (2009). SynFace—speech-driven facial animation for virtual speech-reading support. EURASIP journal on audio, speech, and music processing, 2009, 1–10. doi:https://doi.org/10.1155/2009/191940

Schabus, D., Pucher, M., & Hofer, G. (2013). Joint audiovisual hidden semi-markov model-based speech synthesis. IEEE Journal of Selected Topics in Signal Processing, 8(2), 336–347. doi:https://doi.org/10.1109/JSTSP.2013.2281036

Scott, M. (2007). The Alchemyst: The Secrets of the Immortal Nicholas Flamel. Audiobook.

Sensimetrics. (2014). Speech Test Video Corpus STeVi Retrieved from: https://www.sens.com/products/stevi-speech-test-video-corpus/

Shan, T., & Maddox, R. K. (2021). Speech-in-noise comprehension is improved when viewing a deep-neural-network-generated talking face. Paper presented at the ARO 2021 Mid Winter Meeting. Poster presentation retrieved from https://aro.org/wp-content/uploads/2021/02/Abstract-Book-Cover-Page.pdf

Shinn-Cunningham, B. G., & Best, V. (2008). Selective attention in normal and impaired hearing. Trends in amplification, 12(4), 283–299. doi:https://doi.org/10.1177/1084713808325306

Smayda, K. E., Van Engen, K. J., Maddox, W. T., & Chandrasekaran, B. (2016). Audio-visual and meaningful semantic context enhancements in older and younger adults. PloS one, 11(3), e0152773. doi:https://doi.org/10.1371/journal.pone.0152773

Song, Y., Zhu, J., Li, D., Wang, X., & Qi, H. (2018). Talking face generation by conditional recurrent adversarial network. arXiv preprint 1804.04786. doi:https://doi.org/10.48550/arXiv.1804.04786

Strand, J. F., Brown, V. A., & Barbour, D. L. (2020). Talking points: A modulating circle increases listening effort without improving speech recognition in young adults. Psychonomic Bulletin & Review, 27(3), 536–543. doi:https://doi.org/10.3758/s13423-020-01713-y

Sumby, W. H., & Pollack, I. (1954). Visual contribution to speech intelligibility in noise. The journal of the acoustical society of america, 26(2), 212–215. doi:https://doi.org/10.1121/1.1907309

Suwajanakorn, S., Seitz, S. M., & Kemelmacher-Shlizerman, I. (2017). Synthesizing obama: learning lip sync from audio. ACM Transactions on Graphics (ToG), 36(4), 1–13. doi:https://doi.org/10.1145/3072959.3073640

Tamura, M., Masuko, T., Kobayashi, T., & Tokuda, K. (1998). Visual speech synthesis based on parameter generation from HMM: Speech-driven and text-and-speech-driven approaches. In Proc. Auditory-Visual Speech Processing (pp. 221–224).

Varano, E., Vougioukas, K., Ma, P., Petridis, S., Pantic, M., & Reichenbach, T. (2022). Speech-Driven Facial Animations Improve Speech-in-Noise Comprehension of Humans. Frontiers in Neuroscience, 15. doi:https://doi.org/10.3389/fnins.2021.781196

Vougioukas, K., Petridis, S., & Pantic, M. (2019). End-to-End Speech-Driven Realistic Facial Animation with Temporal GANs. Paper presented at the CVPR Workshops.

Vougioukas, K., Petridis, S., & Pantic, M. (2020). Realistic speech-driven facial animation with gans. International Journal of Computer Vision, 128(5), 1398–1413. doi:https://doi.org/10.1007/s11263-019-01251-8

Yuan, Y., Lleo, Y., Daniel, R., White, A., & Oh, Y. (2021). The Impact of Temporally Coherent Visual Cues on Speech Perception in Complex Auditory Environments. Frontiers in neuroscience, 15, 629. doi:https://doi.org/10.3389/fnins.2021.678029

Yuan, Y., Wayland, R., & Oh, Y. (2020). Visual analog of the acoustic amplitude envelope benefits speech perception in noise. The Journal of the Acoustical Society of America, 147(3), EL246–EL251. doi: https://doi.org/10.1121/10.0000737

